# GESIAP3.0: Sensor-based Image Analysis Program for Transmission Visualization *In Vivo*

**DOI:** 10.1101/2024.10.28.620522

**Authors:** Roger E. Zhu, Xintong Diao, Xiaoyu Liu, Qin Ru, Zenan Wu, Ziyuan Zhang, Loren L. Looger, Jiazhu Zhu

## Abstract

Synaptic transmission mediated by various neurotransmitters influences a wide range of behaviors. However, understanding how neuromodulatory transmitters encode diverse behaviors and affect their functions remains challenging. Here, we introduce GESIAP3.0, an advanced, third-generation image analysis program based on genetically encoded sensors. This tool enables precise quantitative analysis of transmission in both awake, freely moving animals and immobilized subjects. GESIAP3.0 incorporates movement correction algorithms that effectively eliminate image displacement in behaving animals while optimizing synaptic information extraction and simplifying computations on commodity computers. Quantitative analysis of cholinergic, dopaminergic, and serotonergic transmission, corrected for tissue movement, revealed synaptic properties consistent with measurements from *ex vivo* wide-field and *in vivo* two-photon imaging under stable conditions. This validates the applicability of GESIAP3.0 for analyzing synaptic properties of neuromodulatory transmission in behaving animals.

## INTRODUCTION

Intercellular communication in the brain involves both fast-acting transmitters like glutamate and gamma-aminobutyric acid (**GABA**), as well as >500 slow-acting neuromodulatory transmitters such as monoamines, neuropeptides, and other molecules, coordinating diverse behavioral and physiological processes (Greengard, 2001; Sudhof, 2021). Electrophysiological recordings, with their remarkable sensitivity and temporal resolution, have significantly advanced our understanding of fast glutamatergic and GABAergic transmission (von Gersdorff and Borst, 2002; Wu et al., 2014; Pulido and Marty, 2017). The development of high-resolution patch-clamp recordings in the early 1990s revolutionized the quantitative quantal analysis of these synaptic transmissions (Edwards et al., 1990; Stern et al., 1992; Jonas et al., 1993), driving discoveries in silent synapses, synaptic receptor trafficking, synaptic molecular signaling, synaptic nanoscale organization, and other critical insights into brain function and associated disorders (Malinow and Malenka, 2002; Volk et al., 2015; Sudhof, 2021; Connor and Siddiqui, 2023). However, patch-clamp recordings are limited by their inability to capture responses induced by neuromodulators effectively (Dani and Bertrand, 2007; Nadim and Bucher, 2014; Muir et al., 2024). Alternative methods like microdialysis and voltammetry, despite recent advancements, still lack the required spatial or temporal resolution (Olive et al., 2000; Robinson et al., 2008; Darvesh et al., 2011; Li et al., 2022). As a result, while slow-acting neuromodulatory transmitters are implicated in numerous physiological actions and diseases, quantitative and mechanistic insights into their regulation and function remain limited.

The emergence of genetically encoded fluorescence transmitter sensors, many of which emit abundant photons, offers a promising approach to analyzing synaptic transmission (Sabatini and Tian, 2020; Zhu et al., 2020; Labouesse and Patriarchi, 2021; Lin et al., 2021; Wu et al., 2022). Our theoretical framework suggests that advanced image analysis algorithms can harness excess photon signals to enhance spatial and temporal resolution, providing deeper insights into transmission in both healthy and diseased brains (Chen et al., 2021; Lin et al., 2021). Preliminary experiments combining wide-field fluorescence imaging with our genetically encoded sensor-based image analysis programs (**GESIAP1.0** and **GESIAP2.0**) permit quantitative quantal analysis of transmission in *ex vivo* and cultured cell preparations (Zhu et al., 2020; Zheng et al., 2022), validating the idea. These findings have been further validated by high-speed two-photon imaging in intact brains (Kazemipour et al., 2019; Borden et al., 2020). However, two-photon imaging suffers from inefficiencies, low signal-to-noise ratios, and operational complexity. To address these issues, we recently developed the third generation of GESIAP algorithms (**GESIAP3.0**), which, in combination with miniscope-based wide-field imaging, effectively resolves synaptic properties of neuromodulatory transmission in intact brains. GESIAP3.0 integrates movement correction algorithms to eliminate image displacement in both awake and immobilized animals. It also optimizes data extraction from fluorescence responses, simplifying computation to run on an ordinary laptop. Additionally, GESIAP3.0 rescues image sequences acquired under unstable *in vitro* conditions, making synaptic transmission imaging more accessible to less-experienced researchers. Designed to work independently of proprietary software or vendor-specific hardware, GESIAP3.0 allows researchers without programming expertise to analyze neuromodulatory synaptic properties in behaving animals.

## RESULTS

### Development of GESIAP3.0

To monitor synaptic transmission in behaving animals, we developed the third generation of GESIAP algorithms (**GESIAP3.0**) to address image displacement in both awake, freely moving and immobilized animals. We utilized non-rigid movement correction algorithms modified from previous work (Pnevmatikakis and Giovannucci, 2017), focusing on movement correction in the X-Y plane, as wide-field miniscope imaging captures only a single focal plane. Our interactive, piecewise-based algorithm divides frames into smaller, overlapping patches (see Methods), allowing independent adjustments for localized distortions. It generates a mesh of temporal motion vectors to track movement direction and magnitude at each pixel point, adjusting pixel values to compensate for global and local motion. By continuously updating these vectors, the algorithm ensures alignment and correction of movement distortions throughout the image sequence, resulting in a stable, motion-corrected dataset. GESIAP3.0 integrates movement correction with other algorithmic procedures from GESIAP2.0, such as alignment, deconvolution, baseline correction, denoising, background subtraction, and 3D profiling, to process fluorescence image sequences collected from genetically encoded sensor-based functional imaging experiments (**Fig 1A-B**).

**Figure 1.**
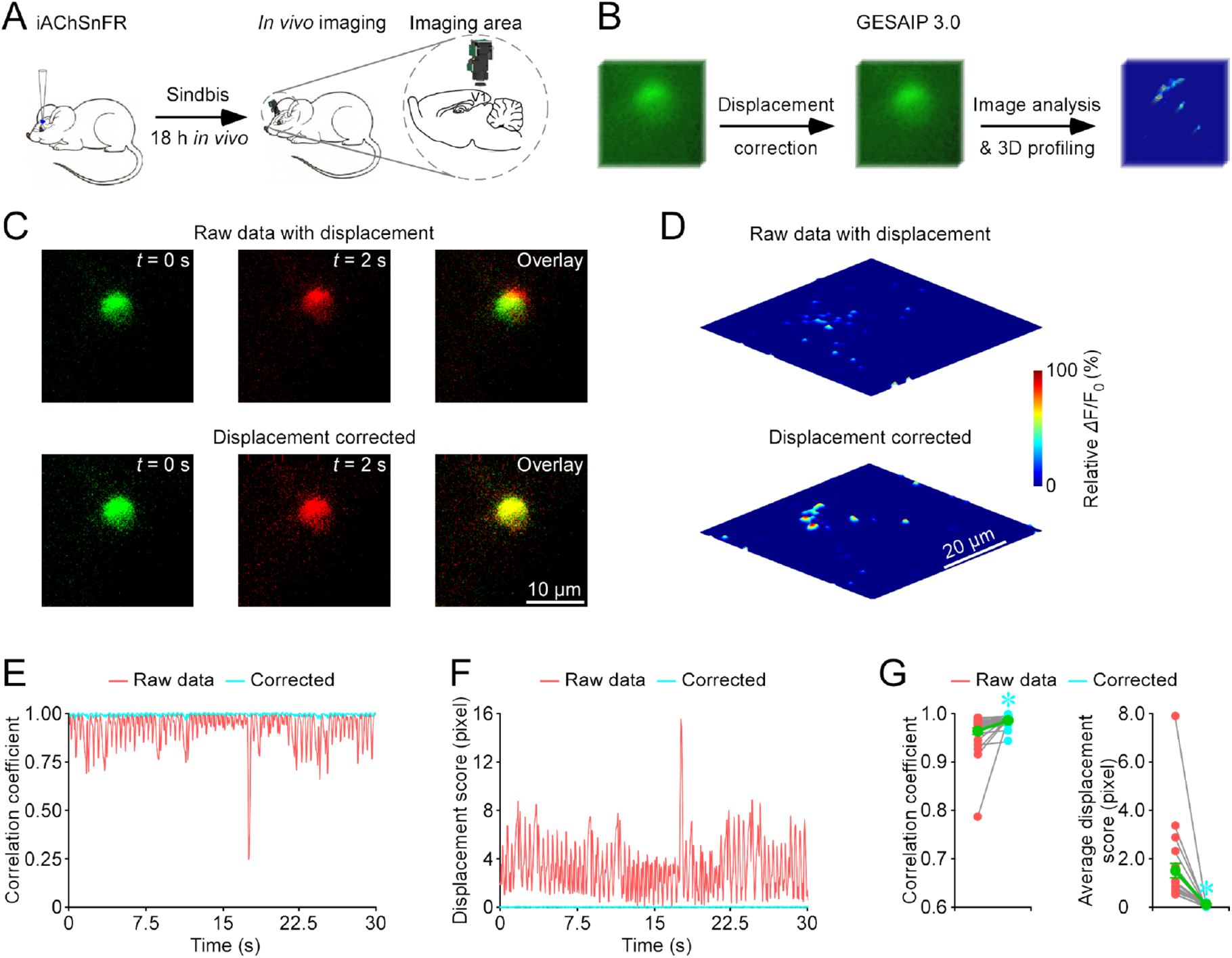
Movement correction algorithms minimize image displacement and improve data fidelity. **(A)** Schematic of miniscope-based imaging of cholinergic responses in the mouse visual cortex *in vivo*. V1: the primary visual cortex. **(B)** GESIAP3.0 incorporates movement correction algorithms alongside procedures for translational alignment, Landweber deconvolution, baseline correction, denoise, background subtraction, *Δ*F/F_0_ heatmap calculation, and three-dimensional (**3D**) transmitter release profiling. **(C)** Snapshots and overlay of images from an iAChSnFR expressing cortical neuron, captured at two time points before and after movement correction. Note the prominent mismatch due to image displacement in the raw data. **(D)** 3D spatiotemporal profiling of electrically evoked fluorescence *Δ*F/F_0_ responses in the iAChSnFR expressing cortical neuron before and after movement correction. Note the absence of prominent peaks in the raw data due to movement displacement. **(E)** Correlation coefficient plots between image frames over time for the iAChSnFR expressing cortical neuron imaged at 16 frames per second. Note the pink and blue traces representing data before and after movement correction, respectively. **(F)** Displacement scores between image frames over time for the same neuron. Note the pink and blue traces representing data before and after movement, respectively. **(G)** Left, average correlation coefficients before and after movement correction (Before: 0.9642±0.0084 vs. After: 0.9872±0.0024; *n* = 25 neurons from 20 animals, *Z* = 4.3733 *p* < 0.0005). Right, average displacement scores before and after movement correction (Before: 1.51±0.30 vs. After: 0.09±0.01; *n* = 25 neurons from 20 animals, *Z* = -4.3733 *p* < 0.0005). Asterisks indicate *p* < 0.05 (Wilcoxon tests).

GESIAP3.0 also features improved denoising algorithms. In GESIAP2.0, a double-exponential synaptic function was used to best fit fluorescence responses (Zheng et al., 2022), which required extensive computation time. To address this, GESIAP3.0 implemented a least-squares non-linear optimization to fit the double-exponential synaptic function using the Levenberg-Marquardt algorithm combining gradient descent with the Gauss-Newton method (Moré, 2006; Gavin, 2019). This approach simultaneously fits both the rise and decay phases of synaptic responses, optimizing parameters for more accurate regression. Additionally, biological constraints based on the response properties of genetically encoded sensors (e.g., (Borden et al., 2020; Sun et al., 2020; Wan et al., 2021)) were incorporated to limit the range of parameter values, enhancing both speed and accuracy. These improvements reduced computation time for analyzing single cell image sequences by ∼10−30-fold, cutting processing time from ∼6−12 hours to ∼20−30 minutes, making analysis feasible on a standard laptop. GESIAP3.0 is hosted on our lab’s GitHub and will be freely accessible to the research community upon publication of the study in a peer-reviewed journal.

### Visualization of synaptic transmission in vivo with GESIAP3.0

We initially tested the applicability of GESIAP3.0 by expressing a genetically encoded sensor for acetylcholine, iAChSnFR (Borden et al., 2020), in the mouse visual cortex using a Sindbis viral expression system (**Fig 1A**). Approximately 24 hours after *in vivo* expression, we implanted a custom-made one-photon fluorescence miniscope (**1P-miniFM**) above the cortex to capture wide-field imaging. To stimulate cholinergic fibers passing through layer I of the cortex (Zhu, 2000; Ray et al., 2014), we positioned a local stimulating electrode on the cortical surface. The evoked fluorescence responses from iAChSnFR-expressing cortical neurons were then imaged using the 1P-miniFM, and GESIAP3.0 was applied to analyze the fluorescence image sequences, enabling us to visualize acetylcholine release (**Fig 1B**).

We found that GESIAP3.0’s movement correction effectively addressed image displacement in the raw image sequences, restoring precise 3D profiling of individual cholinergic releasing synapses that had been previously smeared (**Fig 1C-D**). The performance of the movement correction algorithms was demonstrated by a marked improvement in correlation coefficients, increasing from lows of 0.25−0.50 to an average of 0.98, and a substantial reduction in displacement scores, with pixel displacements decreasing from ∼2 pixels to as low as ∼0.02 pixels post-correction (**Fig 1E-G**).

To further validate the applicability of GESIAP3.0, we analyzed release properties of cholinergic transmission in intact mouse brains. As with previous GESIAP versions (Zhu et al., 2020; Zheng et al., 2022), GESIAP3.0 allowed visualization of acetylcholine diffusion at individual releasing synapses (**Fig 2A**). The movement correction procedure in GESIAP3.0 eliminated spurious signals caused by motion artifacts and sharpened the cholinergic release peaks. Fitting a single-exponential decay function to pixel-wise maximal *Δ*F/F_0_ plots at isolated releasing synapses yielded acetylcholine spatial spread length constants (**Fig 2B-D**). The movement correction decreased measured acetylcholine spatial spread length constants by ∼40%, resulting in a final value of 0.74 µm (**Fig 2E**).

**Figure 2.**
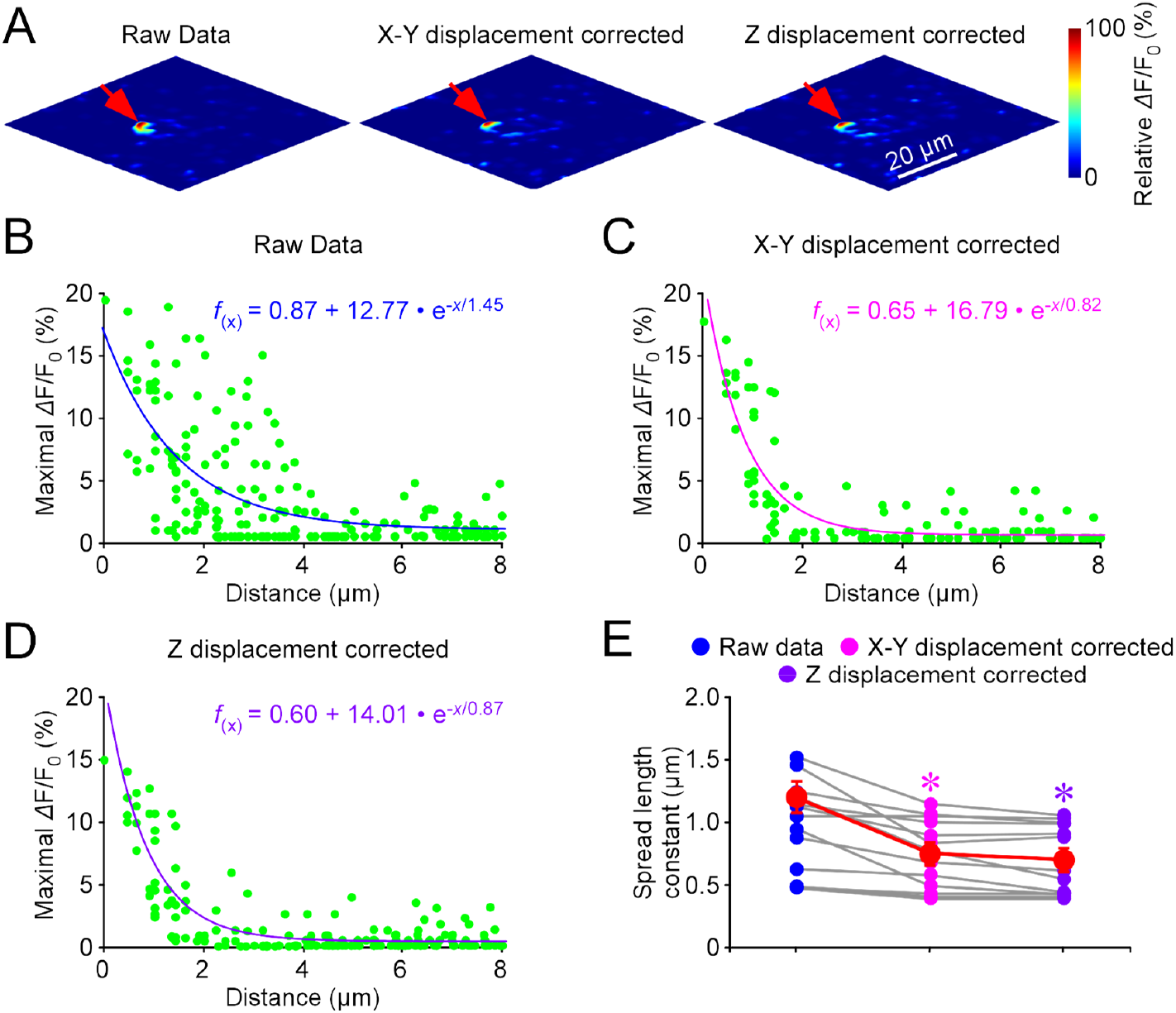
Movement correction algorithms improve image quality. **(A)** 3D spatiotemporal profiling of an iAChSnFR expressing cortical neuron before correction, after movement correction in the X-Y plane, and after additional correction in the Z-axis. **(B)** Pixel-wise maximal *Δ*F/F_0_ at the isolated releasing synapse in the raw image (indicated by a red arrow in **A**). Fitting the data points with a single-exponential decay function (cyan line) yields an estimated ACh spatial spread length constant of 1.45 µm. **(C)** Pixel-wise maximal *Δ*F/F_0_ at the isolated releasing synapse after X-Y plane correction (indicated by a red arrow in **A**). Fitting the data points with a single-exponential decay function (pink line) yields an estimated ACh spatial spread length constant of 0.82 µm. **(D)** Pixel-wise maximal *Δ*F/F_0_ at the isolated releasing synapse after additional Z-axis correction (indicated by a red arrow in **A**). Fitting the data points with a single-exponential decay function (purple line) yields an estimated ACh spatial spread length constant of 0.87 µm. **(E)** Average spatial spread length constants for ACh at cortical neurons from images corrected in the X-Y plane (0.74±0.05 µm; *n* = 10 from 5 neurons, *Z* = -3.059, *p* = 0.002) and after additional Z-axis correction (0.69±0.08 µm; *n* = 10 from 5 neurons, *Z* = -3.189, *p* = 0.001) compared to raw images (1.19±0.10 µm; *n* = 10 from 5 neurons). Note the slightly larger average ACh spread length from X-Y corrected images compared to those with additional Z-axis correction (*Z* = -1.992, *p* = 0.045). Asterisks indicate *p* < 0.05 (Wilcoxon tests).

Next, we enhanced the algorithms to correct for Z-plane drift by compensating for temporal fluctuations within image sequences. Following an initial denoising process, the program identified pixels unlikely to be associated with transmitter releasing synapses or that exhibited peak changes below a defined threshold. The intensity profiles of these non-responsive pixels served as a template to create a Z-plane movement profile, which was then subtracted from each pixel’s temporal trace. This process effectively removed movement noise and corrected for temporal drift while minimizing non-structural artifacts along the Z-plane. Following this correction, a secondary function-fitting denoising cycle was applied, along with additional steps to improve 3D profiling accuracy. Although this method generally improved image sharpness in many cells, it could slightly reduce clarity in others due to the trade-off between signal responses and movement correction (see **Fig 2D** for an example). Overall, Z-plane correction improved acetylcholine spatial spread length constants by approximately 7%, yielding a corrected value of 0.69 µm (**Fig 2E**). This result aligns closely with acetylcholine spatial spread length constants obtained *ex vivo* using wide-field imaging (Zhu et al., 2020; Zheng et al., 2022) and *in vivo* with high-speed two-photon imaging (Kazemipour et al., 2019; Borden et al., 2020), further validating the GESIAP3.0 model.

### Improvement of transmission visualization ex vivo with GESIAP3.0

Despite our *ex vivo* imaging experiments being conducted by experienced scientists with optimized electrophysiology-imaging setups, we encountered tissue movements in a small number of trials, necessitating the exclusion of these results from analysis (Lin et al., 2021). We sought to determine whether GESIAP3.0 could correct for these movements and recover usable data. In a previous study (Zhu et al., 2020), we made Sindbis viral expression of an iAChSnFR in layer 2 stellate neurons of the medial entorhinal cortex in intact mice and then prepared acute entorhinal slices for imaging experiments after ∼18 hours of *in vivo* expression. Tissue movements were observed in some neurons during imaging experiments. By applying GESIAP3.0 to reanalyze these neurons, we observed enhanced sharpness at acetylcholine releasing synapses (**Fig 3A-B**). Additionally, measurements of the acetylcholine spatial spread length constant showed a ∼15% sharpening, reducing the spread length constant from ∼0.85 µm to ∼0.70 µm (**Fig 3C-E**). This latter value aligns closely with those measured from cells without evident movement (Zhu et al., 2020; Zheng et al., 2022). This illustrates the GESIAP3.0’s ability to enhance imaging analysis of neurotransmission in *ex vivo* preparations.

**Figure 3.**
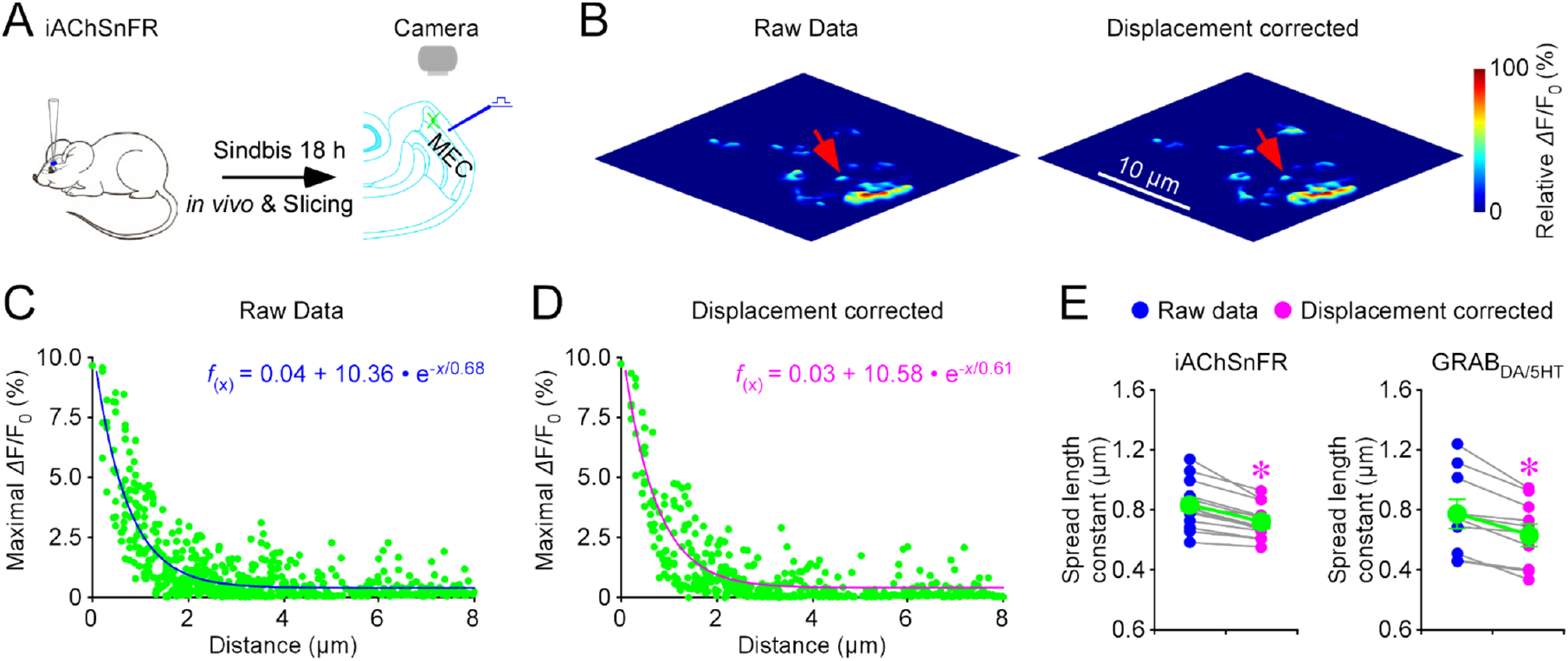
Movement correction algorithms improve image quality in *ex vivo* experiments. **(A)** Schematic of the imaging experiment in a mouse medial entorhinal cortical (**MEC**) slice preparation. **(B)** 3D spatiotemporal profiling of an iAChSnFR expressing entorhinal neuron before and after movement correction in the X-Y plane. **(C)** Pixel-wise maximal *Δ*F/F_0_ at the isolated releasing synapse in the raw image (indicated by a red arrow in **B**). Fitting the data points with a single-exponential decay function (cyan line) yields an estimated acetylcholine spatial spread length constant of 0.68 µm. **(D)** Pixel-wise maximal *Δ*F/F_0_ at the isolated releasing synapse after X-Y plane correction (indicated by a red arrow in **B**). Fitting the data points with a single-exponential decay function (pink line) yields an estimated acetylcholine spatial spread length constant of 0.61 µm. **(E)** Left, average spatial spread length constants for acetylcholine at iAChSnFR expressing entorhinal neurons before and after X-Y plane movement correction (Before: 0.84±0.05 µm; After: 0.72±0.03 µm; *n* = 14 from 6 neurons, *Z* = -3.30, *p* = 0.00012). Right, average spatial spread length constants for dopamine at GABA_DA_ expressing striatal neurons and serotonin at GABA_5HT_ expressing geniculate neurons before and after X-Y plane movement correction (Before: 0.77±0.10 µm; After: 0.63±0.08 µm; *n* = 10 from 4 neurons, *Z* = -2.80, *p* = 0.005). Asterisks indicate *p <* 0.05 (Wilcoxon tests).

In separate sets of experiments from the previous study (Zhu et al., 2020), we used the same *in vivo* Sindbis viral expression to deliver genetically encoded dopamine sensor, GRAB_DA_ (Sun et al., 2020), and serotonin sensor, GRAB_5HT_ (Wan et al., 2021), into the striatum and lateral geniculate nucleus, respectively. We then prepared *ex vivo* thalamic and striatal brain slice preparations to characterize the spatial profile of serotonin and dopamine. Likewise, tissue movements were observed in a few experiments. Applying GESIAP3.0 to reanalyze these image sequences with tissue movements improved sharpness at both dopamine and serotonin releasing synapses (**Fig 3E**). These findings confirm the general applicability of GESIAP3.0 in enhancing imaging analysis of neurotransmission in *ex vivo* preparations.

## DISCUSSION

This study presents GESIAP3.0, the third generation of a genetically encoded sensor-based image analysis program designed for quantitative neurotransmission analysis in behaving animals. GESIAP3.0 features advanced movement correction algorithms that effectively eliminate image displacement artifacts in both freely moving and immobilized animals, as well as *in vitro* preparations. This tool enhances synaptic information extraction and simplifies computational requirements, making it usable on standard laptops without needing proprietary software or vendor-specific hardware. These improvements facilitate more accessible analysis of neuromodulatory synaptic transmission across various contexts, including behavioral studies and *in vitro* or cultured cell preparations.

### General applicability of GESIAP

Tissue displacements caused by voluntary (e.g., licking, chewing, and limb movements), involuntary (e.g., breathing and heartbeats), and reflexive (e.g., muscle contractions and postural shifts) actions present significant challenges for fluorescence imaging and data analysis, particularly in techniques requiring for high spatial or temporal precision (Greenberg and Kerr, 2009; Griffiths et al., 2020). To tackle these issues, we developed GESIAP3.0, an interactive, piecewise, non-rigid correction algorithm program designed to address movement artifacts. Our findings demonstrate that tissue displacements can blur images, but GESIAP3.0 successfully restores image quality.

Our recent studies showed that neuromodulatory transmitters, like acetylcholine, primarily use highly restricted synaptic transmission. Analysis of acetylcholine spatial diffusion in *ex vivo* brain preparations revealed a spread length constant of ∼0.70−0.75 µm (Zhu et al., 2020; Zheng et al., 2022). High-speed two-photon imaging has corroborated these findings, confirming the same spread length constant of ∼0.70−0.75 µm and revealing independent transmitter releases at neighboring releasing synapses in intact brains (Jing et al., 2018; Kazemipour et al., 2019; Borden et al., 2020). By employing GESIAP3.0 for *in vivo* miniscope-based imaging of cholinergic transmission, we identified a similar spread constant, reinforcing the restricted transmission model and validating the utility of GESIAP3.0.

GESIAP3.0 also effectively corrects image displacement in *ex vivo* experiments. While refined electrophysiology-imaging setups and experienced researchers can minimize tissue movement in brain slices, completely preventing displacements remains challenging. Our results show that GESIAP3.0 successfully corrects for these displacements, allowing for data recovery and analysis. Furthermore, the program optimizes synaptic information extraction. It simplifies computations, making it accessible even to researchers with limited experience in high-resolution imaging or access to advanced equipment or computer power. Overall, GESIAP3.0 significantly enhances the toolkit for quantitative analysis of synaptic transmission, with numerous applications in the field.

## METHODS

### Animal preparations

Male and female C57BL/6J mice (Jackson Laboratory, Bar Harbor, ME; RRID: IMSR_JAX:000664), aged 42 to 60 days, were used in this study. Both sexes were included, as sensor-based imaging with genetically encoded sensors showed no differences in fluorescence responses between males and females. Mice were housed in family or pair arrangements at the animal facility of Zhejiang University and the University of Virginia. They were maintained in controlled environments with a temperature of 23±1 °C and humidity of 55±5 %, on a 12-h/12-h light/dark cycle. Food and water were available *ad libitum*. All procedures complied with the guidelines for the Care and Use of Laboratory Animals, as approved by the Laboratory Animal Center of Zhejiang University and the Animal Care & Use Committees of the University of Virginia (Protocol No. 3168), in accordance with US National Institutes of Health guidelines.

### Sindbis preparation and expression

Genetically encoded fluorescent sensors for acetylcholine (iAChSnFR, (Borden et al., 2020)), dopamine, (GABA_DA2m_, (Sun et al., 2020)), and serotonin (GRAB_5HT_, (Wan et al., 2021)) were subcloned into the Sindbis viral construct. Viral particles were produced following established protocols (Jing et al., 2018; Wan et al., 2021). Specifically, fluorescent sensors and their variants were inserted into Sindbis viral vector pSinREP5, using Xba1 and Sph1 restriction enzymes. Neuronal expression was driven by a human synapsin promoter.

Viral expression was performed, as previously reported (Wang et al., 2020; Huang et al., 2021). In brief, mice were anesthetized with sodium pentobarbital at 80 mg/kg or a mix of ketamine and xylazine at doses of 10 mg/kg and 2 mg/kg, respectively, and head-fixed in a stereotaxic frame (RWD Life Science, Shenzhen, China). A craniotomy was created above the injection sites with a 0.5 mm diameter drill bit (RWD Life Science, Shenzhen, China), and ∼50 nl of Sindbis viral solution was pressure-injected into specific brain areas via an injecting micropipette at a rate ∼60 nl/min using a Legato 130 syringe pump (KD Scientific Inc., MA, USA) or a Picospritzer III (Parker Hannifin Corporation, Hollis, NH, USA). The injecting micropipette was held in place for ∼10 minutes post-injection to ensure viral diffusion. Experiments were typically performed within 18±4 hours and 24±4 hours of Sindbis viral infection for *ex vivo* and *in vivo* experiments, respectively.

### Miniscope implementation

Surgical implantation of a home-made one-photon fluorescence miniscope (**1P-miniFM**) was performed following a previously described protocol (Larkum et al., 2001). Mice were initially anesthetized with 3% isoflurane (0.5 l/min) and maintained with 1.5% isoflurane (0.1 l/min). Pressure points and incision sites were sterilized with 75% ethanol and infiltrated with 0.25% lidocaine. Body temperature was maintained at 37.2 ± 0.3°C. After head-fixation, a ∼4 mm diameter hole was drilled into the skull above the targeted cortical regions, with bone debris flushed out using artificial cerebrospinal fluid (**ACSF**) (in mM: 124 NaCl, 3 KCl, 1.25 NaH2PO4, 26 NaHCO3, 2 MgCl2, 2 CaCl2, pH 7.4). The dura mater was carefully removed, and the cortical surface was kept moist with ACSF.

The miniscope was positioned above the exposed cortex and adjusted for optimal visualization of sensor expressing cells. A concentric bipolar stimulating electrode (CE2C65; FHC, Bowdoin, ME) was placed 100-500 µm from the target cells to evoke neurotransmitter release (Cat. #30209, FHC, ME, USA). Single electric stimulating pulses (200 ms, up to 20 V) were generated by a stimulator (Master 8, A.M.P.I., Tektronix, Israel) and delivered via a stimulation isolator (ISO-Flex, A.M.P.I., Tektronix, Israel). Imaging was performed either under continuous anesthesia or in awake, immobilized animals.

### Fluorescence imaging

Wide-field epifluorescence *in vivo* imaging was performed using the 1P-miniFM, which integrates an electrowetting (fine) lens and a motorized mechanical (coarse) focusing systems. Fluorescent sensor expressing cells in intact brains were excited by a 460−480-nm LED (LXZ1-PB01; Lumileds, San Jose, CA, USA), collimated with a molded aspheric lens (#83-605; Edmund, Santa Monica, CA, USA), and filtered through a light filter set consisting of ET470/40x, ET525/50m, and T495lpxr. Fluorescence signals were captured by an AR1335CSSM11SMD20 sensor (Onsemi, Pheonix, AZ, USA), featuring advanced 1.1 µm pixel size BSI technology, and sampled with a 16-bit A/D converter.

Wide-field epifluorescence *in vitro* imaging was performed using a Hamamatsu ORCA FLASH4.0 camera (Hamamatsu Photonics, Shizuka, Japan). Fluorescent sensor expressing cells in acutely prepared tissue slices were excited by a 460-nm ultrahigh-power low-noise LED (Prizmatix, Givat-Shmuel, Israel) (Wang et al., 2015; Jing et al., 2018), typically set at 0.20 mW/mm^2^ unless stated otherwise. The FLASH4.0 camera operated at a frame rate of 10-50 Hz, using an Olympus 40x water-immersion objective with a numerical aperture of 0.8.

The microscopic point-spread functions were measured with 100 nm green beads FluoSpheres (F8803, Invitrogen, CA, USA) (**Fig S1**). Despite variability in basal fluorescence F_0_ across the entire expressing neuron surfaces due to heterogeneous iAChSnFR and GRAB_DA_ sensor expression, *Δ*F/F_0_ responses showed no or weak correlation with F_0_, suggesting independence from sensor expression levels and reliability in measuring transmitter release (**Fig S2**) (cf. (Zhu et al., 2020; Zheng et al., 2022)).

### Movement correction with GESIAP3.0

To address brain movement in behaving animals, we employed non-rigid movement correction algorithms adapted from previous work (Pnevmatikakis and Giovannucci, 2017). Our focus was on correcting movement in the X-Y plane using a piecewise-based NoRMCorre algorithm. Unlike rigid methods that only manage uniform translations and rotations, NoRMCorre enhances movement tracking by dividing the field of view into smaller, overlapping patches. In our approach, frames were split into 32×32 pixel patches with an 8-pixel overlap. This configuration allows the algorithm to adjust each patch independently, accounting for localized, non-uniform distortions instead of applying a uniform correction across the entire image. The overlapping nature of the patches ensures smoother transitions and more precise corrections across different regions. To achieve sub-pixel accuracy, the algorithm calculates shifts for each patch, generating a mesh of temporal motion vectors. These vectors adjust the pixel values, ensuring they move smoothly across frames without altering the actual pixel positions. It constructs an initial median template from the first 200 frames, which is then updated every 200 frames by averaging the most recently registered frames. This continuous adaptation process improves motion correction by refining the alignment of patches. As the vectors are updated throughout the image sequence, the algorithm effectively corrects movement distortions, resulting in a stable, motion-corrected dataset.

We calculated correlation coefficients and pixel displacements between consecutive frames to evaluate motion correction effectiveness. Higher correlation coefficients indicate better alignment, while smaller pixel displacements reflect successful artifact reduction. Iterative implementation of the algorithm proved unnecessary, as negligible differences were observed in most cells, with slight degradation in a few others.

We also developed an algorithm to correct Z-plane movement. This algorithm identified significant peaks related to stimuli following a denoising step, categorizing background pixels with peak changes below a predefined threshold. Background traces were averaged and subtracted from individual pixel traces to correct for Z-plane displacements. The corrected datasets underwent a second iteration of function-fitting denoising, followed by additional procedures to finish 3D profiling.

### Imaging analysis with GESIAP3.0

After movement correction, we analyzed fluorescence responses using the updated GESIAP3.0, developed in MATLAB 2023a with the Image Processing Toolbox and Curve Fitter (Mathworks). GESIAP3.0 comprises five major algorithmic procedures, including alignment, deconvolution, baseline adjustment, denoising, and background subtraction. The alignment procedure utilizes translational intensity-based automatic registration (Reddy and Chatterji, 1996), further minimizing average pixel displacement through a non-parametric diffeomorphic image registration algorithm (Vercauteren et al., 2009).

The subsequent deconvolution corrects light distortions using empirically obtained point spread functions (Zhu et al., 2020) and Landweber algorithms, which outperformed various other methods, including naïve/regularized inverse filtering, Tikhonov regularization, Tikhonov-Miller, and Richardson-Lucy algorithms (Zheng et al., 2022).

By integrating our pre- and post-processing procedures, Landweber algorithms achieved performance comparable to commercial packages like Huygens (Ponti et al., 2007). This deconvolution process iteratively minimizes a least-squares cost function while imposing non-negativity constraints (Landweber, 1951). We found that 25-50 iterations typically balanced image enhancement and noise amplification, with the latter largely mitigated by the denoising step.

Baseline adjustments compensated for small fluorescence decay due to photobleaching by applying double-exponential fits to the first and last 10 seconds of recordings. Individual pixel adjustments produced new flat baselines for the entire image sequence. Since genetically encoded transmitter sensors behave similarly to transmitter receptors in binding transmitters, we tested three model synaptic functions — simple rise and decaying, alpha (Rall, 1967), and double-exponential (Destexhe et al., 1994) — to fit fluorescence responses, ultimately selecting the double-exponential model due to its superior performance. The response onset is determined by back-extrapolation of the linear line crossing 40% and 80% response points in the rising phase. A bilateral non-linear filter (Tomasi and Manduchi, 1998) was applied as needed to further reduce noise before synaptic function fitting. Background subtraction adjusted levels by uniformly subtracting an average fluorescence value from a non-responsive region adjacent to the cell, thereby improving image presentation without inflating *Δ*F/F_0_ values.

### Synaptic transmission property analysis with GESIAP3.0

To visualize individual transmitter releasing synapses and estimate postsynaptic transmitter spatial diffusion extent, the maximal electrically evoked maximal *Δ*F/F_0_ responses at individual pixels over time were plotted to create 3D spatial profiles for individual transmitter releasing synapses. Individual releasing synapses were isolated and identified using the density-based spatial clustering algorithm DBSCAN (Ester et al., 1996). Iterative selection and analysis isolated and identified individual releasing synapses from overlapping clusters. Pixels with the maximal *Δ*F/F_0_ responses in individual releasing synapses were assumed to be the centers of release. Using strategies from the super-resolution localization microscopy analysis (Thompson et al., 2002; Sauer, 2013; Small and Stahlheber, 2014), fluorescence *Δ*F/F_0_ intensity profiles were averaged over multiple exposures, multiple releases and/or multiple directions of transmitter diffusion gradients, and fit with a single-exponential decay function. Fitting was performed at well-isolated releasing synapses, and their decay constants were extracted as spatial spread length constants.

### Statistical analysis

Statistical results were reported as mean±s.e.m. Animals were randomly assigned into control or experimental groups, investigators were blinded to experiment conditions, and no sample was excluded for analysis. Given the negative correlation between the variation and square root of sample number, *n*, the group sample size was typically set to be ∼8−25 to optimize the efficiency and power of statistical tests. Statistical significances of the means (*p*<0.05; two sided) were determined using Mann-Whitney Rank Sum non-parametric tests, and statistical significances of the linear relationships of two data groups were determined using linear regression *t* tests. The normal distribution and similar variance within each comparison group of data were checked prior to statistical tests. The data that support the findings of this study are available from the corresponding author upon request.

## ACKNOWLEDGMENTS

We thank members of the Ke Si and Julius Zhu labs for suggestions and technical support.

## AUTHOR CONTRIBUTIONS

REZ and JZZ conceived the concept and led the project; REZ developed MATLAB-based image analysis program and analyzed data with assistance from XD, XL, QR, ZW, ZZ, and JZZ; XD and XL performed biology experiments; LLL provided key reagents; REZ wrote the manuscript with input from all other coauthors.

## FIGURE LEGENDS

**Figure S1.**
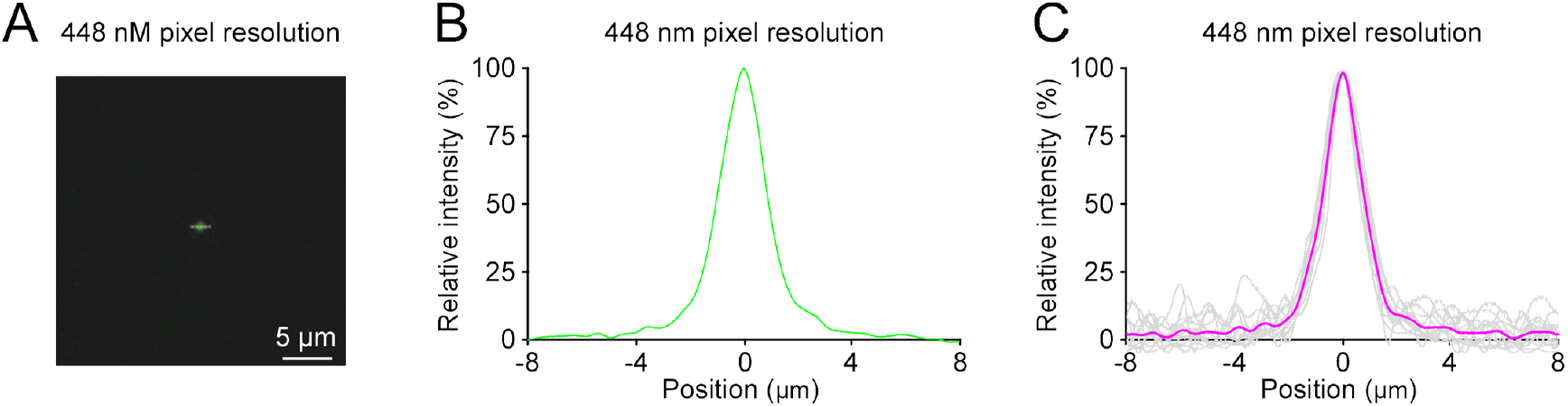
Point-spread function (PSF) determination for the miniscope. **(A)** Fluorescence image of a 100-nm green GATTA bead acquired using a miniscope at 448 nm pixel resolution. **(B)** Corresponding point-spread function (**PSF**) of the 100-nm green GATTA bead shown in (**A**) obtained using the miniscope at 448 nm pixel resolution. **(C)** Individual (light gray) and average (pink) PSFs of 100-nm green GATTA beads obtained under the miniscope at 448 nm pixel resolution. Note the full width at half maximums (**FWHMs**) of PSFs to be 1.63±0.08 μm (*n* = 13).

**Figure S2.**
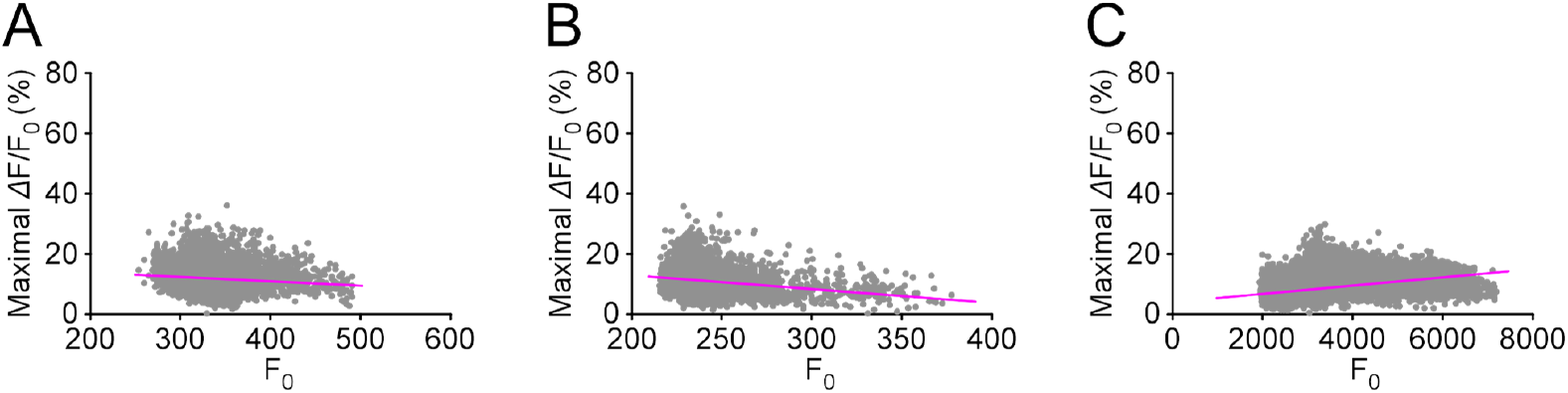
Fluorescence responses are largely independent of sensor expression levels. **(A)** Plot of ΔF/F_0_ against F_0_ of the iAChSnFR expressing mouse cortical neuron shown in Figure 1 (Slope of regression line = -0.000151; Normality test *p* ≤ 0.001; Constant variance test *p* ≤ 0.001; *r*^2^ = 0.09048; *F* = 91.09; *n* = 10,201; *p* < 0.0005; Linear regression *t* test). **(B)** Plot of ΔF/F_0_ against F_0_ of the iAChSnFR expressing mouse cortical neuron shown in Figure 2 (Slope of regression line = -0.000466; Normality test *p* ≤ 0.001; Constant variance test *p* ≤ 0.001; *r*^2^ = 0.1942; *F* = 399.68; *n* = 10,201; *p* ≤ 0.001; Linear regression *t* test). **(C)** Plot of ΔF/F_0_ against F_0_ of the iAChSnFR expressing mouse accumbal neuron shown in Figure 3 (Slope of regression line = 0.000014; Normality test *p* ≤ 0.001; Constant variance test *p* ≤ 0.001; *r*^2^ = 0.1797; *F* = 2233.87; *n* = 10,201; *p* < 0.0005; Linear regression *t* test).

